# Air-Liquid Interface Model for Influenza Aerosol Exposure In Vitro

**DOI:** 10.1101/2024.11.04.621830

**Authors:** Brittany Seibert, C. Joaquin Caceres, L. Claire Gay, Nishit Shetty, Flavio Cargnin Faccin, Silvia Carnaccini, Matthew Walters, Linsey C. Marr, Anice C. Lowen, Daniela S. Rajao, Daniel R. Perez

## Abstract

Airborne transmission is an essential mode of infection and spread of influenza viruses among humans. However, most studies use liquid inoculum for virus infection. To better replicate natural airborne infections *in vitro*, we generated a calm-aerosol settling chamber system designed to examine the aerosol infectivity of influenza viruses in different cell types. Aerosol inoculation was characterized for multiple influenza A virus (FLUAV) subtypes, including a pandemic 2009 H1N1, a seasonal swine H3N2, and an avian H9N2 using this exposure system. While each FLUAV strain displayed high infectivity within MDCK cells via liquid inoculation, differences in infectivity were observed during airborne inoculation. This was further observed in recently developed immortalized differentiated human airway epithelial cells (BCi-NS1.1) cultured in an air-liquid interface. The airborne infectious dose 50 for each virus was based on the exposure dose per well. Our findings indicate that this system has the potential to enhance our understanding of the factors influencing influenza transmission via the airborne route. This could be invaluable for conducting risk assessments, potentially reducing the reliance on extensive and costly *in vivo* animal studies.

**Importance:** This study presents a significant advancement in influenza research by developing a novel in vitro system to assess aerosol infectivity, a crucial aspect of influenza transmission. The system’s ability to differentiate between mammalian-adapted and avian-adapted influenza viruses based on their aerosol infectivity offers a valuable tool for pre-screening the pandemic potential of different strains. This could potentially streamline the risk assessment process and inform public health preparedness strategies. Moreover, the system’s capacity to examine aerosol infectivity in human airway epithelial cells provides a more relevant model for studying virus-host interactions in natural airborne infections. Overall, this study provides an accessible platform for investigating aerosol infectivity, which could significantly contribute to our understanding of influenza transmission and pandemic preparedness.

## Introduction

Influenza A virus (FLUAV) has caused recurrent epidemics associated with severe disease and zoonotic transmission from animals to humans (1). FLUAV results in seasonal epidemics in the global population, partially controlled by annual vaccination; however, occasional zoonotic transmission of FLUAV can potentially lead to a pandemic outbreak affecting millions of individuals worldwide. Sporadic human infections with H5, H7, and H9 avian influenza viruses have been reported (2). Further, bidirectional human-to-swine transmission of H1 and H3 FLUAVs has been documented for several decades (3). Generally, zoonotic events are restricted to sporadic primary individual cases; however, FLUAVs have the potential to acquire the ability to transmit between mammals, which is a critical step leading towards the start of a pandemic (1). Therefore, understanding the processes involved in FLUAV transmission between hosts, particularly airborne transmission, can contribute to pandemic risk assessment and public health preparedness.

Current literature indicates two primary modes of FLUAV transmission: contact between an infected and a susceptible individual (direct or indirect) or airborne transmission (1, 4, 5). Airborne transmission occurs through the inhalation of infectious virus carried in respiratory aerosol particles as large as 100 µm. While the density of infectious respiratory particles is more concentrated closer to the source, larger-sized particles are relevant only at a close range due to their inability to travel greater distances before falling to the ground within 1-2 m of the infectious host. Meanwhile, smaller-sized particles are relevant at both close and long ranges, as they remain suspended in the air for an extended period of time and travel many meters away from the source. While a range of particle sizes are detected from respiratory fluids expelled from an infected individual, the deposition site of a particle also depends on the release (breathing or exhaled puff cloud) and environmental factors, including air velocity, temperature and relative humidity. However, when examining infectious respiratory particles, previous studies reported higher FLUAV vRNA loads in smaller particles compared to larger infectious respiratory particles (1, 4–16)

While airborne transmission is a standard route of FLUAV infection in humans, many *in vivo* experiments are conducted via liquid or direct inoculation. Previous studies examined differences in pathogenesis when inoculating animal models via intranasal or aerosol (17–22). Conclusively, aerosol inoculation in mice, ferrets, and non-human primates had little effect on the overall clinical outcome but more closely resembled FLUAV infection in humans, including slower disease progression and increased homogeneity of pulmonary lesions (17, 20–22). While animal models, principally ferrets, recapitulate human disease regarding susceptibility, pathogenesis, and transmission, ferrets are expensive and potentially difficult to work with in high numbers (23). Therefore, assessing aerosol infectivity in an *in vitro* system can be valuable for screening viruses for potentially enhanced human-to-human transmission before or in conjunction with *in vivo* studies.

Infection of adherent cell monolayers with a liquid inoculum to study virus replication is an established method, allowing for testing of multiple experimental conditions and cell lines of different origins with considerable reproducibility. However, traditional liquid infection for *in vitro* replication poorly represents natural exposure and infection of cells within the respiratory tract (24). A previous study adapted a multi-component aerosolized system to inoculate epithelial cell monolayers with high pathogenicity avian influenza (HPAI) H5N1, a low pathogenicity avian influenza (LPAI) H7N9, and a seasonal H3N2 FLUAV (24). While the previous aerosol system incorporates numerous environmental components and considerations, we aimed to develop a calm-aerosol settling chamber system that is accessible and cost-effective for analyzing aerosol infection efficiency of different FLUAV viruses *in vitro*. Utilizing this aerosol system, we successfully infected a common adherent monolayer cell line, Madin-Darby canine kidney (MDCK) cells, and differentiated human airway epithelial cells cultured in air-liquid interface (ALI) conditions with three FLUAV subtypes: a pandemic H1N1, swine-origin H3N2, and avian-origin H9N2. The aerosol infectious dose 50 (AID_50_) for each virus and cell type was calculated by determining the cumulative amount of virus deposited into a cell culture plate within the total time of exposure, considering a downward flux of particles at a defined settling velocity reflective of the aerosol exposure system. The aerosol exposure system resulted in infectious virus replication in two different cell lines, suggesting that this system can be utilized to investigate aerosol infectivity of circulating FLUAVs *in vitro* prior to expensive animal experimentation studies.

## MATERIALS AND METHODS

### Cells

MDCK cells were a kind gift from Robert Webster (St Jude Children’s Research Hospital, Memphis, TN, USA). Cells were maintained in Dulbecco’s Modified Eagles Medium (DMEM, Sigma-Aldrich, St Louis, MO, USA) containing 10% fetal bovine serum (FBS, Sigma-Aldrich, St. Louis, MO, USA), 1% antibiotic/antimycotic (AB, VWR, Radnor, PA, USA) and 1% L-Glutamine (Sigma-Aldrich, St Louis, MO, USA). Cells were cultured at 37°C under 5% CO2.

Human airway epithelial cells BCi-NS1.1(25) were obtained from Dr. Ronald Crystal (Weill Cornell Medicine, NY, USA). Cells were maintained in 1X Basal Media containing PneumaCult-Ex Plus Basal Medium (490 mL) (STEMCELL Technologies, Vancouver, Canada), PneumaCult-Ex Plus 50X Supplement (STEMCELL Technologies, Vancouver, Canada), 0.1% Hydrocortisone (STEMCELL Technologies, Vancouver, Canada), 1% Penicillin-Streptomycin (5000 U/mL; ThermoFisher Scientific, MA, USA), 0.5% Amphotericin B (ThermoFisher Scientific, MA, USA), and 0.5% Gentamycin (50 mg/mL; Sigma-Aldrich, St Louis, MO, USA). Cells were cultured at 37°C under 5% CO2. Trypsin-EDTA (Sigma-Aldrich, ST Louis, MO, USA) followed by a solution of 85% HEPES Buffered Saline (Lonza, Basel, Switzerland) with 15% FBS (Sigma-Aldrich, St. Louis, MO, USA) were used for passaging the cells. The BCi-NS1.1 cells were not used after passage 25. Differentiation of the BCi-NS1.1 was performed in 12 mm trans-well plates with 0.4 µm pore polyester membrane inserts (Corning Inc., NY, USA). Before plating, trans-well membranes were coated with human type IV collagen (Sigma-Aldrich, St. Louis, MO, USA) and rinsed with 1X phosphate-buffered saline (PBS; ThermoFisher Scientific, MA, USA). Once dried, 300,000 BCi-NS1.1 cells/well were plated into the trans-well membrane insert with 1X Basal Media and cultured at 37°C under 8% CO2. After reaching 100% confluency, the cells were changed to air-liquid interface (ALI) conditions by removing the apical media and then changing the basal media for 1X ALI media. ALI media contained PneumaCult ALI Base Medium (450 mL) (STEMCELL Technologies, Vancouver, Canada), 10X PneumaCult ALI Supplement (50 mL) (STEMCELL Technologies, Vancouver, Canada), 1% Penicillin-Streptomycin (5000 U/mL; ThermoFisher Scientific, MA, USA), 0.5% Amphotericin B (ThermoFisher Scientific, MA, USA), 0.5% Gentamycin (50 mg/mL; Sigma-Aldrich, St Louis, MO, USA), 1% PneumaCult ALI Maintenance Supplement (STEMCELL Technologies, Vancouver, Canada), 0.2% Heparin solution (STEMCELL Technologies, Vancouver, Canada), and 0.5% Hydrocortisone Stock solution (STEMCELL Technologies, Vancouver, Canada). Cells were cultured in ALI conditions at 37°C under 8% CO2 for 5 days and then incubated at 37°C under 5% CO2 until they reached 17 days in ALI conditions.

### Viruses

Virus stocks of the mouse-adapted H1N1 A/California/04/2009 (Ca04; H1N1), swine-origin A/turkey/Ohio/313053/04 (Oh/04; H3N2), and avian-origin A/Guinea Fowl/Hong Kong/WF10/99 (WF10; H9N2) were generated by reverse genetics as previously described (26). Virus stocks were amplified in 10-day-old specific pathogen-free (SPF) embryonated chicken eggs. Virus stocks were titrated by tissue culture infectious dose 50 (TCID_50_), and virus titers were established by the Reed and Muench method (27). Virus sequences were confirmed by next-generation sequencing as previously described (28).

### Histology and Immunofluorescence of differentiated HAE cells

Differentiation of HAE cells in ALI culture were confirmed using immunofluorescence staining of paraffin-embedded cross-sections or by top-staining of the trans-well membrane. Briefly, trans-well membranes were cut out of the trans-well, rolled into a cassette, and fixed in 10% neutral-buffered formalin (NBF) for at least 72 h. The cells were then embedded in paraffin and processed for routine histopathology with hematoxylin and eosin staining (HE). Non-stained slides were used for immunofluorescent staining. Slides were de-paraffinized in xylene and ethanol as previously described (29). Heat antigen retrieval was performed by steaming the slides for 45 min with a citrate buffer solution, followed by a PBS wash. For top staining, the trans-wells were fixed directly with 4% paraformaldehyde in PBS for 30 min and then washed 2X with PBS. Samples were then permeabilized with 0.3% triton X-100 in PBS, followed by blocking with 5% bovine serum albumin in PBS for 1 h at room temperature. The samples were then stained with the following primary antibodies: KRT5 (basal cell; 2 µg/mL; HPA059479; Millipore Sigma, MA, USA), beta-tubulin I and II (cilia; 1/800; T8535, Sigma-Aldrich, MO, USA), CC16 (Club cell; 2 µg/mL; RD181022220-01; BioVendor, NC, USA), MUC5AC (M1 mucin; 2 µg/mL; MA5-12178, ThermoFisher, MA, USA), and TFF3 (goblet cell; 2 µg/mL; HPA035464; Millipore Sigma, MA, USA) for 1 hr at room temperature. To visualize the staining, samples were incubated with Alexa Fluor 488 Goat Anti-Mouse IgG (dilution 1/1000; A-11032, ThermoFisher, MA, USA) or Alexa Fluor 594 Donkey Anti-Rabbit IgG (dilution 1/1000; A-21207; ThermoFisher, MA, USA) as secondary antibodies with DAPI to identify cell nuclei for 1 h at room temperature. Actin staining was performed by incubating samples with ActinGreen 488 ReadyProbes (R37110, ThermoFisher, MA, USA) for 5 min. Subsequently, membranes were mounted using Vectashield plus antifade mounting medium (Vector laboratories, CA, USA). Immunofluorescent microscope images were collected using a Nikon confocal microscope with a 60x lens. Images were captured and adjusted using Nikon NIS-Elements software (v 4.60.00).

For ⍺2,3 and ⍺2,6 receptor staining, samples were not permeabilized until after the secondary antibody incubation. Cells were stained with the following primary antibodies: Maackia Amurensis Lectin II (⍺2,3, Biotinylated, 15 µg/mL; B-1265-1; Vector laboratories, CA, USA) and Sambucus Nigra Lectin (⍺2,6, Fluorescein, 15 µg/mL; F13012; Vector Laboratories, CA, USA) for 30 min at room temperature. The samples were then incubated with secondary Streptavidin Alexa Fluor 594 conjugate (1/1000; S32356; ThermoFisher, MA, USA) for 30 min at room temperature. Cells were then permeabilized for 10 min and stained with DAPI for 10 min to visualize the cell nuclei. The membranes were mounted using Vectashield plus antifade mounting medium and imaged. Integrated density was calculated from fluorescent confocal images using ImageJ (30) and divided by the number of nuclei. Statistical analysis was performed using a paired t-test. Receptor distribution was analyzed in triplicates.

### Aerosol exposure design

The bioaerosol system developed was adapted from previously published designs (20, 24, 31). Included within the aerosol chamber setup is an air pump (flow rate 13.3 L/min), Buxco mass dosing controller (Data Sciences International, MN, USA), Aeroneb lab control module (Kent Scientific, CT, USA), Aeroneb lab nebulizer unit (Small VMD; Kent Scientific, CT, USA), Mass dosing exposure chamber (Data Sciences International, MN, USA), SKC BioSampler (SKC, PA, USA), BioLite+ High-volume sample pump (SKC, PA, USA), SKC vacuum pump (SKC, PA, USA), liquid traps (Fisher Scientific, NH, USA), HEPA-CAP filters (VWR, PA, USA), and multiple sizes of plastic tubing and tubing adapters. The particle size distribution generated by the Aeroneb lab nebulizer loaded with 6 mL of PBS was measured using an Aerotrak 9306 particle counter (TSI, MN, USA). The Aerotrak instrument counts particles in six bins with minimum sizes of 0.6, 1, 2, 4, 6 and 10 µm, and it was programmed to sample every 30 seconds for 15 min. The Aeroneb lab nebulizer unit was connected to the Aeroneb lab control module, which was then connected to the Buxco Mass dosing controller to control the nebulizer output efficiency and the duration (min) of aerosol generation. An input of HEPA-filtered air into the mass dosing chamber was provided by pooled air from the air pump and Buxco mass dosing controller for a combined flow rate of 16 L/min. Simultaneously, the air was sampled from the exposure chamber via the SKC BioSampler using the BioLite+ High-volume sample pump at 12.5 L/min. In a second port, virus-laden aerosols were pulled through a liquid trap and a HEPA-CAP filter by the SKC vacuum pump at 3.5 L/min. Prior to each exposure, the flow rates for each component were checked using a 4100 series flowmeter (TSI, MN, USA).

### Biosampler collection from aerosol exposure chamber

To examine the amount of aerosolized virus in the chamber, two biosamplers were included within the chamber setup: the SKC BioSampler and the NIOSH biosampler. For each experiment, 6 mL of 1 × 10^6^ TCID_50_/mL of each virus was aerosolized using the Aeroneb lab nebulizer as previously described. Aerosol sample collection was performed for 15 min. The SKC BioSampler contained 10 mL of Opti-AB and ran at a rate of 12.5 L/min, while the NIOSH biosampler comprised a 15 mL conical tube, 1.5 mL screw-cap tube, and PTFE filter (0.3 µm), each sampling particles in a different size range, that ran at a rate of 3.5 L/min. After collection, an aliquot of the SKC BioSampler was stored at -80°C for future titration. For the NIOSH biosampler, 4 mL of Opti-AB was used to wash the first-stage tube, 1.5 mL of Opti-AB was used to wash the second-stage tube, and 4 mL of Opti-AB was used to wash the filter after aerosol collection. Media collected/used for washing the NIOSH were stored at -80°C for future titration. The liquid media collected from the SKC BioSampler and NIOSH was titrated for infectious virus by TCID_50_ or vRNA quantification by a real-time reverse transcriptase PCR (RT-qPCR). The concentration of infectious virus and vRNA in the aerosols (C_aer_) collected by each biosampler was calculated by multiplying the liquid media volume (V_samp_) by the concentration of virus in it (C_samp_) and dividing by the volume of air sampled, which is the product of the time of exposure (t) and the sampler’s air flow rate (Q_samp_), as previously described (20):

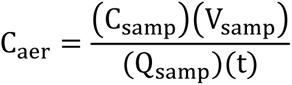

### RNA extraction and RT-qPCR

RNA was extracted from the biosampler collection using the MagMax-96 AI/ND viral RNA isolation kit (ThermoFisher Scientific, MA, USA) following the manufacturer’s protocol. Primers M+25 (AGATGAGTCTTCTAACCGAGGTCG) and M-124 (TGCAAAAACATCTTCAAGTCTCTG) were used for titrating samples inoculated with H9N2 (WF10). Primer M-124_Ca/04_CJC (TGCAAAGACACTTTCCAGTCTCTG) and M-124_Ty/04_CJC (TGCAAAAACGTCTTCGAGTCTCTG) were used in replacement of M-124 for H1N1 (Ca04) and H3N2 (Oh/04) vRNA titration, respectively. A probe with FAM as a reporter and TAMRA as a quencher was used (56-FAM/TCA GGC CCC CTC AAA GCC GA/36-TAMSp). The RT-qPCR was performed using a Quantabio qScript XLT One-Step RT-qPCR ToughMix kit (Quantabio, MA, USA) in a 20 µL final reaction volume on the QuantStudio 3 Real-Time PCR System (ThermoFisher Scientific, MA, USA). Each reaction contained 1X master mix, 0.5 µM of each primer, 0.3 µM probe, and 5 µL of RNA. The RT-qPCR cycling conditions were 50°C, 20 min; 95°C, 1 min, 40 cycles at 95°C, 1 min; 60°C, 1 min; and 72°C 1s; with a final cooling step at 4°C. A standard curve was generated using 10-fold serial dilutions of the virus stock of a known titer to correlate RT-qPCR crossing point (Cp) values with the viral load. RNA virus loads were calculated as log_10_ TCID_50_ equivalents/mL.

### Virus studies in vitro: Direct Inoculation versus Aerosol Exposure

Confluent (90%) monolayers of MDCK cells within a 6-well plate were directly inoculated with 500 µL of ten-fold dilutions of a multiplicity of infection (MOI) starting at 0.01 (1.1 – 1.96 × 10^3^ TCID_50_/mL) until 10^−6^ for each virus. Plates were incubated for 1 hr and rocked every 15 min at 37°C under 5% CO2. The virus inoculum was removed, and the cells were washed twice with 1 mL of PBS. Subsequently, 2 mL of Opti-AB containing 1 μg/mL N-p-tosyl-ʟ-phenylalanine chloromethyl ketone (TPCK)-treated trypsin (Worthington Biochemicals, NJ, USA) was added to each well. For aerosol exposure, Opti-AB (500 µL/well) was added to the MDCK cells before aerosol exposure to prevent drying and cell death. After the 15-min aerosol exposure, cells were removed from the chamber and incubated for 1 hr at 37°C under 5% CO2. The 500 µL of Opti-AB added before exposure was removed, and the cells were washed twice with 1 mL of PBS. Subsequently, 2 mL of Opti-AB containing 1 μg/mL TPCK-treated trypsin was added to each well.

At 0-, 12-, 24-, 48-, and 72 hours post-inoculation (hpi), 200 µL of tissue culture supernatant was collected (replaced with an equal volume of Opti-AB containing 1 μg/mL TPCK-treated trypsin) and 50 µL of the volume collected was used for a hemagglutination assay (HA) described below. The rest of the volume (150 µL) was stored at -80°C for infectious virus quantification by TCID_50_ following the Reed and Muench method (27). For each experiment, 6 wells of cells were inoculated per dilution, and each experiment was performed twice.

Differentiated HAE cells incubated in ALI conditions for 17 days were washed 3 times with PBS before inoculation to remove accumulated mucus. Cells were directly inoculated with 500 µL of ten-fold dilutions starting at an MOI 1 (3.41 – 3.71 × 10^6^ TCID_50_/mL) until 0.01 for each virus. For aerosol exposure, the same virus dilutions were used and the differentiated HAE cells were exposed to aerosolized virus for 15 min. After aerosol exposure, cells were removed from the chamber. Directly inoculated and aerosol exposed cells were incubated for 1 hr at 37°C under 5% CO_2_. Negative controls included direct inoculation or aerosolization of 500 µL Opti-AB. Subsequently, the virus inoculum was removed, the cells were washed with 1 mL of PBS three times, and 1.5 mL of new ALI media was added to the basal compartment. At the indicated time points, 200 µL of Opti-AB was added to the apical side of the well, and then cells were incubated for 10 min at 37°C under 5% CO_2_. The media was then collected, and 50 µL was used for HA while the rest was stored at -80°C for infectious virus quantification by TCID_50_ assay (27). For each experiment, 6 wells of cells from a 12 well transwell plate were inoculated per dilution, and each experiment was performed twice.

### Hemagglutination Assay (HA)

Hemagglutination units (HAU) were calculated as previously described (32). Diluted RBCs (turkey for H1N1 and H3N2 and chicken for H9N2) were incubated with two-fold serial dilutions of cell culture supernatant. For MDCK experiments, HAU were determined after 45 min incubation at room temperature. For differentiated HAE cells, HAU were determined after overnight incubation at 4°C as previously described for differentiated nasal epithelial cells (32).

### Calculation of the aerosol infectious dose 50

Utilizing different parameters within the aerosol exposure chamber system design, we calculated the cumulative virus deposition per 6-well plates over the total 15-min exposure (**Supplemental Fig 1**). The cumulative virus deposition was based on the expected inoculum. The proportion of wells with detectable infectious virus and their respective dilution factor/virus deposition was analyzed in a non-linear regression model. To determine the aerosol infectious dose 50 (AID_50_), we calculated the half-maximal effective concentration (EC_50_) for each experiment.

### Graphs/Statistical Analyses

Data analyses and graphs were performed using GraphPad Prism software version 9.4 (GraphPad Software Inc., CA, USA). Statistical analysis when comparing viruses was performed using a two-way ANOVA with Tukey correction. A P value below 0.05 was considered significant.

## RESULTS

### Aerosol exposure system

We generated a calm-aerosol settling chamber system that is accessible and cost-effective for analyzing aerosol infection efficiency of different FLUAV viruses (**Fig 1**). HEPA-filtered air was inputted into the exposure chamber through a pooled source of air outputted from the Buxco Mass dosing chamber (3-5 L/min) and a commercial air pump (13.3 L/min) for a combined rate of 16 L/min after filtration. The Buxco mass dosing controller was also utilized to regulate the output efficiency and duration (min) of aerosol generation of the nebulizer using the Aeroneb lab control module. Aerosols were generated using the Aeroneb lab nebulizer small VMD unit that expels smaller particles between 2.5 - 4 µm. Evaluation of the aerosol particle distribution from the small VMD nebulizer showed that the majority of the expelled particles were identified to be 0.6 – 2 µm in size (**Fig 2A**). An average of 48.78% (mean (x̄): 2.5 × 10^6^ particles) of the collected aerosols were classified to be 0.6 µm, 43.78% (x̄: 2.3 × 10^6^ particles) classified as 1 µm, 7.43% (x̄: 4.1 × 10^5^ particles) as 2 µm, 0.001% (x̄: 51 particles) as 4 µm, and less than 0.0001% (x̄: 0.23 particles) 6 µm and (x̄: 0.07 particles) 10 µm. To examine the concentration of infectious virus in expelled aerosols, samplers including the SKC BioSampler (12 L/min) and the NIOSH biosampler (3.5 L/min) coupled with the BioLite+ High-volume sample pump (12 L/min) and SKC vacuum pump (3.5 L/min) were utilized. The SKC BioSampler has been commonly used to collect infectious FLUAV (33), while the NIOSH biosampler can segregate aerosols of different sizes (first stage: >4 µm, second stage: 1-4 µm, third stage: <1 µm) (31). Viable viral particles expelled from the Aeroneb nebulizer from a liquid inoculum were examined for infectious virus and vRNA when 1 × 10^6^ TCID_50_/mL of three different FLUAV subtypes, including a pandemic H1N1 (Ca04), swine-origin H3N2 (Oh/04), and avian-origin H9N2 (WF10), was aerosolized. Infectious virus titers (**Fig 2B; bars**) and vRNA (**Fig 2B; circles**) are shown in the concentration of infectious virus and vRNA in the aerosols (C_aer_; TCID50/L of air), as collected by the SKC BioSampler (20). Infectious virus titers and vRNA were similar among all three viruses collected in the SKC BioSampler, suggesting similar viability post-aerosolization of all three viruses. Utilizing the NIOSH biosampler, vRNA loads were categorized based on size; however, infectious virus was not examined as it was below the detection limit (data not shown). While similar vRNA loads were detected in the inoculums (2-way ANOVA; p > 0.05), lower vRNA loads were observed for H1N1 compared to H3N2 in the first and third stage (2-way ANOVA; first stage: p=0.04, second stage: p= 0.07, third stage: p=0.005) and H9N2 (2-way ANOVA; first stage: p<0.001, second stage: p= 0.007, third stage: p<0.001) in all stages (**Fig 2C**). While the first two stages were similar among H3N2 and H9N2, H3N2 had significantly lower vRNA loads than H9N2 (2-way ANOVA: p<0.001) in the last stage of the NIOSH. In conclusion, infectious virus and vRNA were detected among both biosamplers, validating the production of infectious virus-laden particles from the Aeroneb nebulizer utilized in this system. Interestingly, differences in vRNA loads among the different particle sizes were observed among the FLUAV subtypes.

**Figure 1.**
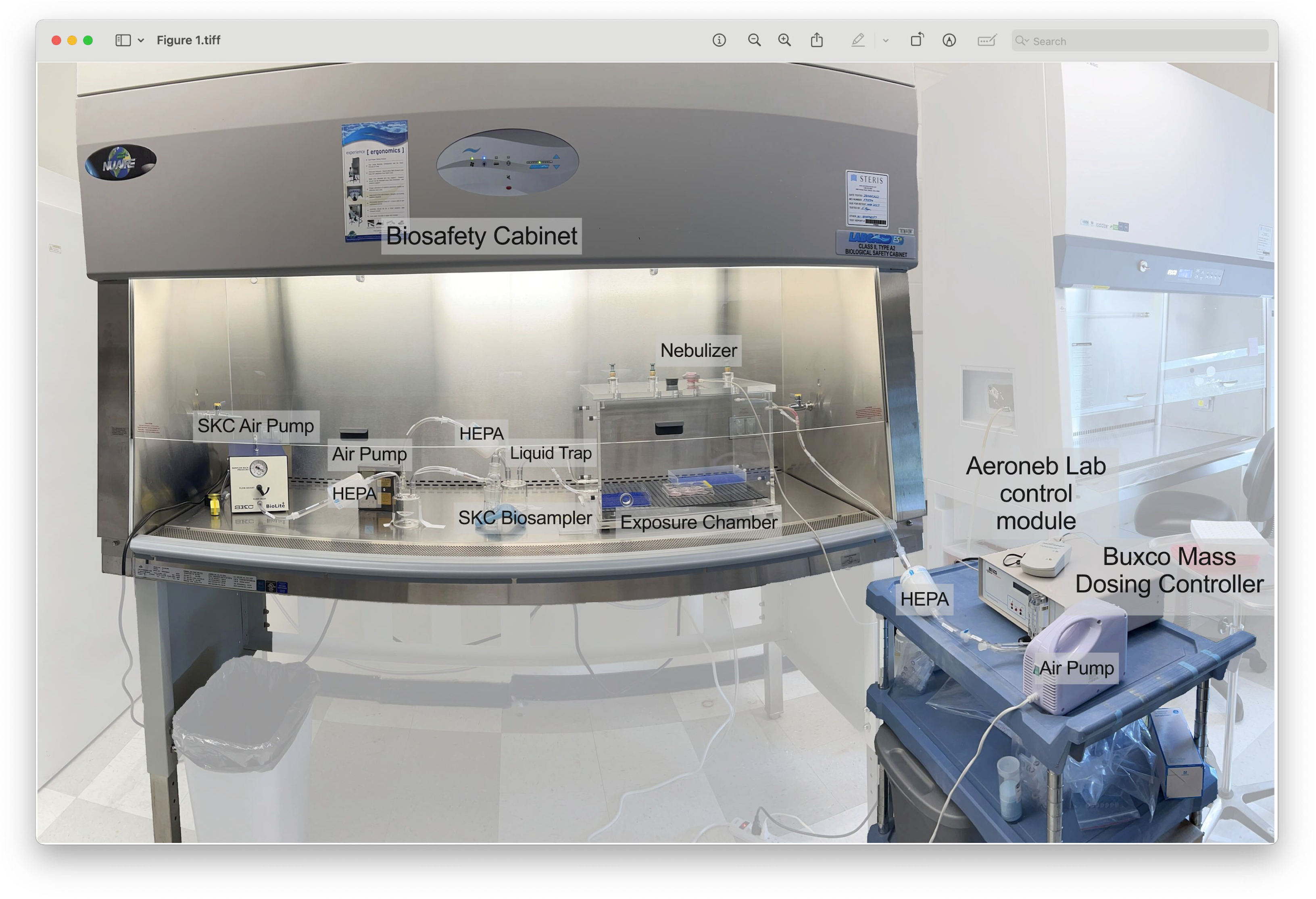
Experimental Aerosol System Set Up. The aerosol exposure system consists of an air pump (flow rate 13.3 L/min), the Buxco mass dosing controller, Aeroneb lab control module, Aeroneb lab nebulizer unit, the Mass dosing exposure chamber, SKC BioSampler, BioLite+ High-volume sample pump, SKC vacuum pump, liquid traps, HEPA-CAP filters and multiples sizes of plastic tubing and tubing adapters. FLUAV was aerosolized directly into the exposure chamber using the Aeroneb nebulizer. Virus-laden aerosols were collected from one output port into the SKC BioSampler, followed by a liquid trap and HEPA filter before entering a vacuum pump. In a second output port, tubes are connected to a liquid trap and HEPA filter before entering a vacuum pump. Each experiment consisted of a 15-min exposure followed by a 5-min purge before removing the cells from the exposure chamber. Cells were placed in the same orientation and location for each exposure.

**Figure 2.**
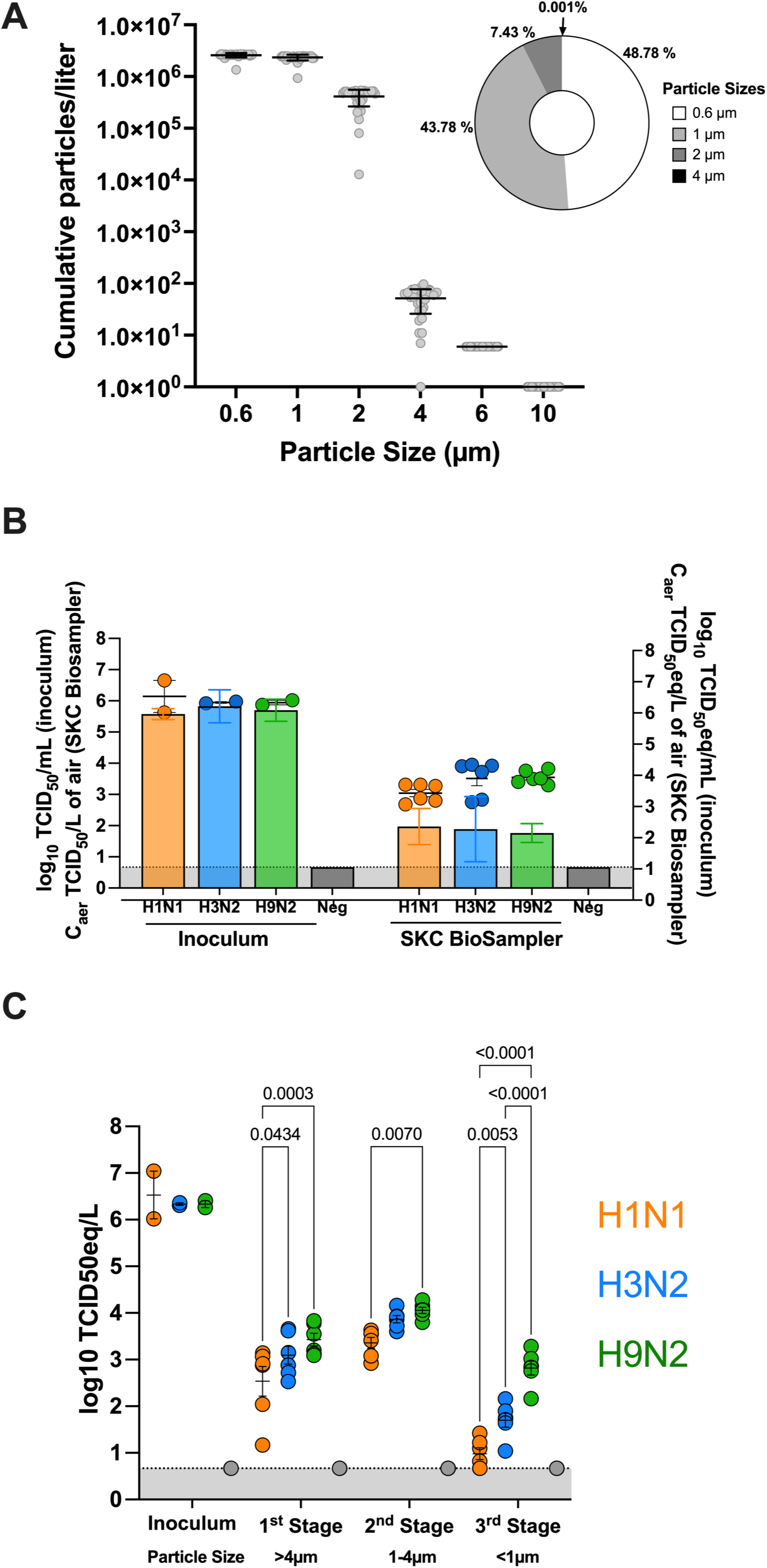
Characterization of the *in vitro* aerosol exposure system and biosamplers. (**A**) The Aerotrak particle counter was used to evaluate the particle size distribution generated by the Aeroneb nebulizer utilized in the aerosol exposure design. Aerosols were measured every 30 seconds for a total of 15 min. The cumulative particle size is illustrated in a bar graph. The percentage of particles that were classified as 0.6, 1, 2, or 4 µm are illustrated in a pie chart. Percentages for 6 µm and 10 µm are not shown since the percentage is <0.0001%. The size indicates the lower end of the size range. **(B)** Bars represent infectious virus from samples collected from the SKC BioSampler (left axis) were quantified and expressed as concentration of infectious virus in the aerosols (C_aer_; TCID50/L of air). Dots represent vRNA loads (right axis) were quantified by RT-qPCR and calculated as a TCID_50_ equivalent (C_aer_; TCID50eq/L of air). Back titrations of virus inoculum are expressed as log_10_ TCID_50_/mL (left axis). vRNA loads of virus inoculum are expressed as log_10_ TCID50eq/mL (right axis). Statistical analysis was performed using a two-way ANOVA with Tukey correction. **(C)** vRNA load quantification from samples collected from each stage of the NIOSH biosampler are displayed as a TCID_50_ equivalent (C_aer_; TCID50eq/L of air). Viruses are distinguished by color (H1N1: orange, H3N2: blue, H9N2: green). Statistical analysis was performed using a two-way ANOVA with Tukey correction.

### Aerosol infection of FLUAV subtypes H1N1, H3N2 and H9N2 in MDCK cells

MDCK cells are commonly utilized in FLUAV research and have been previously demonstrated to support the replication of numerous FLUAV subtypes (34). Therefore, MDCK cells were exposed via aerosol (AR) to ten-fold dilutions of H1N1, H3N2 or H9N2 starting at MOI of 1 × 10^−2^ TCID50/well for 15 min, followed by a 5 min purge to enable the clearing of the aerosolized virus from the chamber. Mock infections using PBS led to no visible decrease in cell adherence. For comparison purposes, cells were directly inoculated via liquid of the same inoculum and volume for each virus. Assessed by hemagglutination assay (HAU > 2) for infectious virus post-aerosol-inoculation at 24 hpi, H1N1 (**Fig 3A**, n=10/12 wells) had the greatest number of wells at an MOI 1 × 10^−2^ TCID50/well while H3N2 and H9N2 had no wells above the limit of detection (**Fig 3B and C**, respectively). Several wells with detectable HAU were observed at 48 hpi and 72 hpi via aerosol and liquid inoculation. Therefore, cell supernatant collected at 48 h after aerosol and liquid inoculation of each virus was titrated for infectious virus by TCID_50_ **(Fig 4)**. For all three viruses, a greater proportion of wells were positive when virus was delivered in a liquid inoculum compared to aerosol inoculation. When aerosolized, H1N1 (**Fig 4A**) had the greatest number of positive wells (5/12, 41.6%) after aerosol inoculation at an MOI 1 × 10^−4^ TCID50/well, followed by H3N2 and H9N2 (**Fig 4B and C**; 3/12, 25.0%) with an equal number of positive wells at the same MOI. However, the viral titers reached in positive wells were more variable for aerosol inoculation but did not show significant differences between aerosol and liquid inoculation.

**Figure 3.**
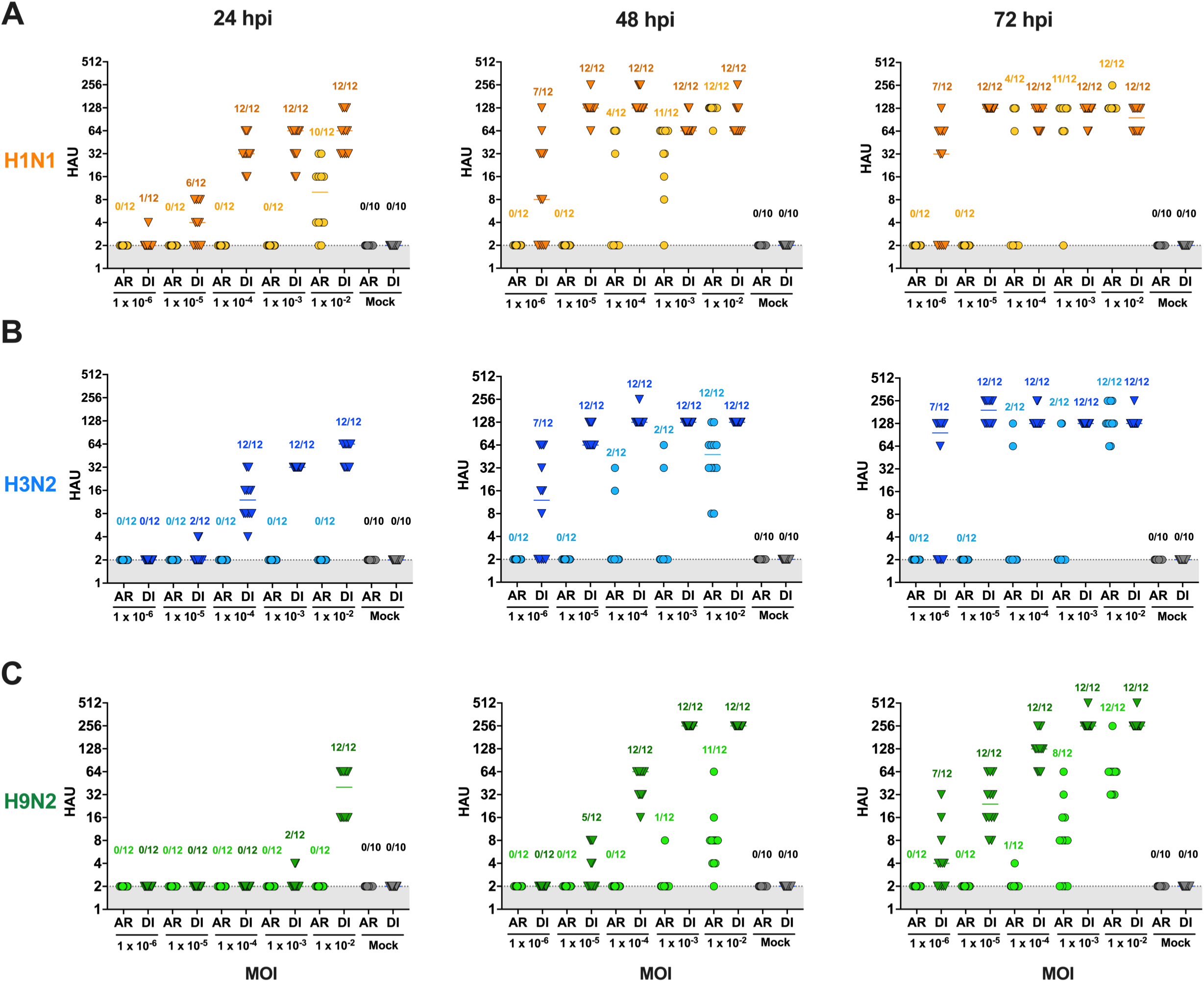
Hemagglutination (HA) assay titration of H1N1, H3N2, and H9N2 FLUAV post aerosol and direct inoculation in MDCK cells. MDCK cells were inoculated via aerosol (AR; circles) or direct (DI; triangles) inoculation of ten-fold dilutions from 1 MOI of **(A)** H1N1, **(B)** H3N2, and **(C)** H9N2. Cell culture supernatant was collected at 0-, 12-, 24-, 48-, and 72 hours post-inoculation (hpi). The first two time points (0 and 12 hpi) resulted in HAU below the limit of detection and are therefore not shown. Each point represents that HAU from 1 well. Numbers above the points represent the number of wells considered positive/total number of wells. Wells with HAU > 2 were considered positive. Viruses are distinguished by differences in color (Ca04: orange, Oh/04: blue, WF10: green).

**Figure 4.**
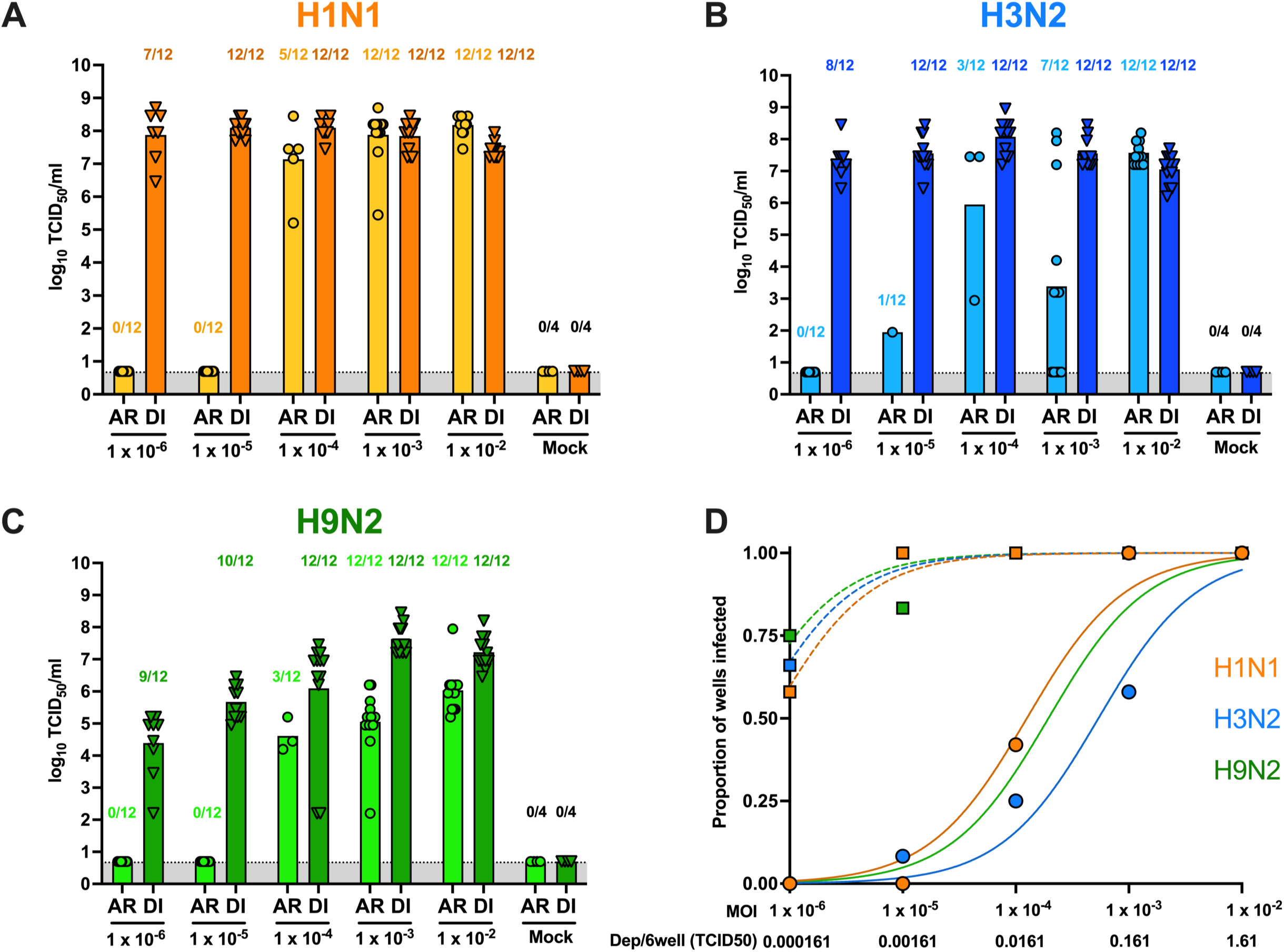
FLUAV exposure in MDCK cells via aerosol or direct inoculation. MDCK cells were exposed to 10-fold dilutions of (**A**) H1N1, (**B**) H3N2, or (**C**) H9N2 from 1 × 10^−6^ to 1 × 10^−2^ MOI via aerosol (AR) or direct inoculation (DI). Cell supernatant was collected at 48 hours post-inoculation (hpi) and infectious virus was quantified and expressed as log10 TCID_50_/mL. Numbers above each bar represent the number of wells considered positive/total number of wells. Mean titers were calculated on based on number of positive wells, which equals to levels of virus replication achieved when infection was productive **(D)** The proportion of wells with detectable infectious virus observed in A, B, and C and their respective dilution factor was analyzed in a non-linear regression model to calculate the half maximal effective concentration (EC50). The cumulative deposition of aerosol/6 well plate for each inoculum is displayed below. Solid lines with circles indicate aerosol exposure, while dashed lines with squares represent liquid inoculation. Viruses are distinguished by color (H1N1: orange, H3N2: blue, H9N2: green).

Utilizing the number of positive wells at each MOI dilution after direct and aerosol inoculation, we calculated the aerosol infectious dose 50 (AID_50_) for each virus in MDCK cells (**Fig 4D**). The estimated cumulative deposition of aerosolized virus/6-well plate was calculated incorporating the average inoculum of all runs (MOI 0.01 x̄: 1.33 × 10^3^ TCID_50_/mL) with aerosol system parameters including volume nebulized, exposure time, nebulizer flow rate, volume of the chamber, well area, flow rate of the 5 min purge, and deposition velocity of an aerosol particle with a size of 2.5-4 µm (**Supplemental Figure 1**). The AID_50_ was determined using a non-linear regression model of the proportion of positive wells by TCID_50_ and their respective dilution to calculate the half-maximal effective concentration (EC_50_). After aerosol inoculation, H1N1 had the lowest AID_50,_ with 0.021 TCID_50_ cumulative aerosol deposition/6 well plate (MOI = 1 × 10^−3.9^ TCID50/well), followed by H9N2 at 0.035 (MOI: 1 × 10^−3.7^ TCID50/well), and H3N2 at 0.13 TCID_50_ (MOI: 1 × 10^−3.3^ TCID50/well)) (**Fig 4D, circles**). Similar EC50s were observed for all three viruses for directly inoculated MDCK cells (MOI: H1N1 = 10^−6.1^, H3N2 = 10^−6.3^, H9N2 = 10^−6.4^) (**Fig 4D, squares**). The AID_50_ results demonstrate different aerosol infection efficiencies among the three viruses, particularly between H1N1 and H3N2, in MDCK cells despite similar liquid inoculation efficiency and replication.

### FLUAV infection in differentiated human airway epithelial cells BCi-NS1.1

We next investigated the ability to infect a more relevant system using differentiated human airway epithelial cells (HAE) via aerosol inoculation. For this, we used BCi-NS1.1 cells, previously immortalized by infecting human airway basal cells obtained from brushings of the airway epithelium of a healthy nonsmoker with a retrovirus expressing human telomerase (25). Immortalizations of the BCi-NS1.1 cells enable multipotent differentiation capacity for over 40 passages, resulting in numerous *in vitro* advantages; however, little is known about the replication capacity of the BCi-NS1.1 cells for different subtypes of FLUAV. Therefore, we investigated FLUAV sialic acid receptor distribution and replication kinetics of a pandemic H1N1, swine-origin H3N2, and avian-origin H9N2 FLUAV. Non-differentiated BCi-NS1.1 cells showed a limited number of cell layers resembling a monolayer phenotype (**Fig 5A**, **left**), whereas differentiated BCi-NS1.1 cells incubated in ALI conditions for 17 days displayed characteristics of pseudostratified ciliated epithelium with visible cilia with intermittent presence of goblet cells (**Fig 5A, right**). The presence of ciliated cells (β-tubulin I and II) at the top of the epithelium and basal cells (KRT-5) located closer to the membrane were confirmed by immunofluorescent staining (**Fig 5B**). Club cells (CC16), goblet cells (TFF3), and mucin (MUC5AC) were also confirmed in the differentiated BCI-NS1.1 cells (**Fig 5C**), further supporting the conclusion that BCi-NS1.1 cells incubated in ALI conditions for 17 days resemble a pseudostratified ciliated HAE. The distribution of FLUAV cell receptors within the differentiated BCi-NS1.1 cell line has not been reported. Therefore, we examined the presence of ⍺2,3- and ⍺2,6-sialic acids (SA) in 17-day ALI cultures. Hemagglutinin from human-origin FLUAV preferentially binds α2,6-SA while hemagglutinin from avian-origin FLUAV preferentially binds to α2,3-SA (35). Differentiated BCi-NS1.1 showed fluorescence positivity for both ⍺ 2,3- and ⍺ 2,6-SA; however, there was significantly more integrated density/number of nuclei and visible staining of the α2,6-SA compared to the α2,3-SA (**Fig 5D**). We next examined virus replication of H1N1, H3N2, and H9N2 in BCi-NS1.1 cells via direct inoculation. While the virus replication kinetics were similar between H1N1 and H3N2, H9N2 had limited virus replication that decreased until 72 hours post-inoculation (hpi) (**Fig 5E**), suggesting restrictive replication of the avian-origin H9N2 FLUAV within the differentiated BCi-NS1.1 cells.

**Figure 5.**
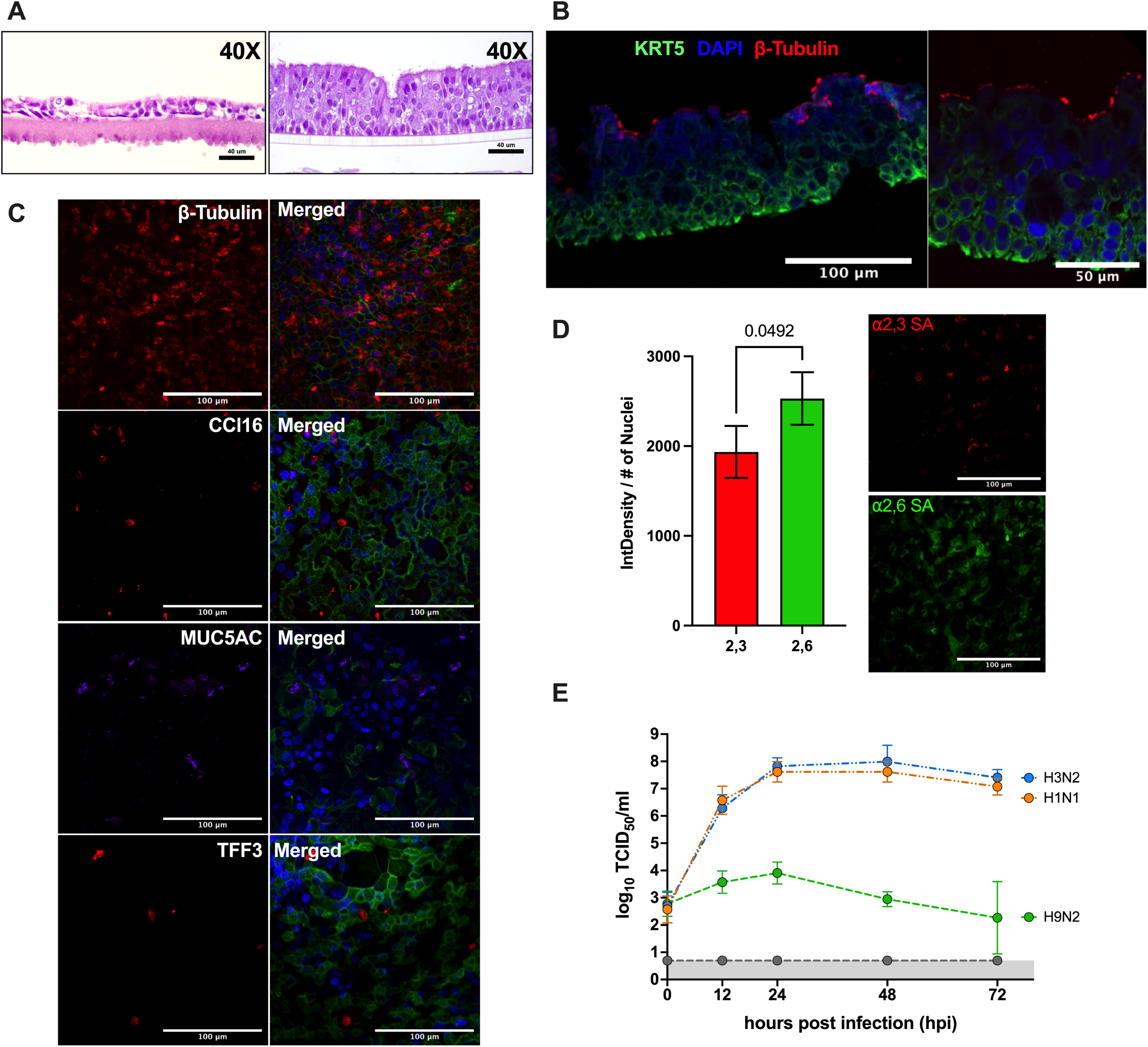
Characterization and FLUAV replication in differentiated BCi-NS1.1 cells. BCi-NS1.1 cells were cultured under air-liquid interface (ALI) conditions for 17 days. **(A)** Paraffin-embedded cross sections of BCi-NS1.1 cells prior to ALI conditions (left) and after 17 days in ALI (right) with H&E staining. Images were captured at 40X magnification. **(B)** Paraffin-embedded cross sections of ALI cultured BCi-NS1.1 cells stained by immunofluorescence with KRT5 (basal cell marker; green), DAPI (cell nuceli; blue), and β-tubulin I and II (ciliated cell marker; red). Images were captured using Nikon confocal microscope with NIS Elements software at 40X (left) and 60X (right). **(C)** ALI cultured BCi-NS1.1 cells were fixed with 4% paraformaldehyde and stained for β-tubulin I and II (ciliated cell marker; red), CC16 (club cell marker; red), MUC5AC (mucin makrer), TFF3 (goblet cell marker), actin (green), and DAPI (cell nuclei; blue). Each panel with separate staining from the same column was merged in the last row. Images were captured using Nikon confocal microscope with NIS Elements software at 60X. **(D)** ALI-cultured BCi-NS1.1 cells were fixed with 4% paraformaldehyde and stained for ⍺ 2,3-(red) and ⍺ 2,6-(green) sialic acids (SA) distribution. Integrated density was calculated from fluorescent confocal images using ImageJ and divided by the number of nuclei. Statistical analysis was performed using a paired t-test. **(E)** ALI-cultured BCi-NS1.1 cells were infected with 1 MOI of H1N1, H3N2, and H9N2. Samples were collected from the apical part of the trans-wells at 0-, 12-, 24-, 48-, and 72 hours post-infection. The samples were titrated by TCID_50_. Viruses are distinguished by differences in color.

### Aerosol infection of FLUAV in differentiated human airway epithelial cells BCi-NS1.1

Since the H1N1 and H3N2 strains replicated efficiently in the differentiated BCi-NS1.1 cells, we investigated the ability to infect the HAE cells via aerosol using the exposure system developed. Differentiated BCi.NS1.1 cells were inoculated with different MOIs (0.01, 0.1 and 1) of H1N1 and H3N2 utilizing the aerosol exposure system described previously. A subset of cells was directly inoculated via liquid with the same inoculum for each virus. Early time points in infection (12 and 24 hpi) resulted in all wells below 2 HAU for direct and aerosol inoculations and were therefore not further examined. For both viruses, a larger proportion of wells were positive when virus was delivered in a liquid inoculum compared to aerosol inoculation at 48 and 72 hpi (**Figure 6**). Post aerosol-inoculation, detectable HAU for H1N1 and H3N2 virus strains were observed at all three MOIs. HAU were below the detection limit at MOIs of 10 to 0.01 at all time points for differentiated BCi-NS1.1 cells exposed to H9N2 via aerosol (data not shown). Cell supernatant titrated for infectious H1N1 and H3N2 virus were similar among the three MOIs (**Fig 7A** and **7B**). Next, we calculated the AID_50_ for each virus in the differentiated HAE cells by calculating the EC50. The average inoculum of all runs (MOI 1 x̄: 3.56 × 10^6^ TCID50/mL) was incorporated to calculate the average deposition per 6-transwells over the 15-min exposure. H3N2 had the lowest AID_50_ at 66.2 TCID_50_ cumulative aerosol deposition/6 well plate (MOI: 1 × 10^−1.8^), followed by H1N1 at 105.2 TCID_50_ cumulative aerosol deposition/6 well plate (MOI: 1 × 10^−1.6^) (**Fig 7C**). Taken together, the AID_50_ results suggest successful aerosol inoculation of a pandemic H1N1 and swine-origin H3N2 in differentiated HAE cells.

**Figure 6.**
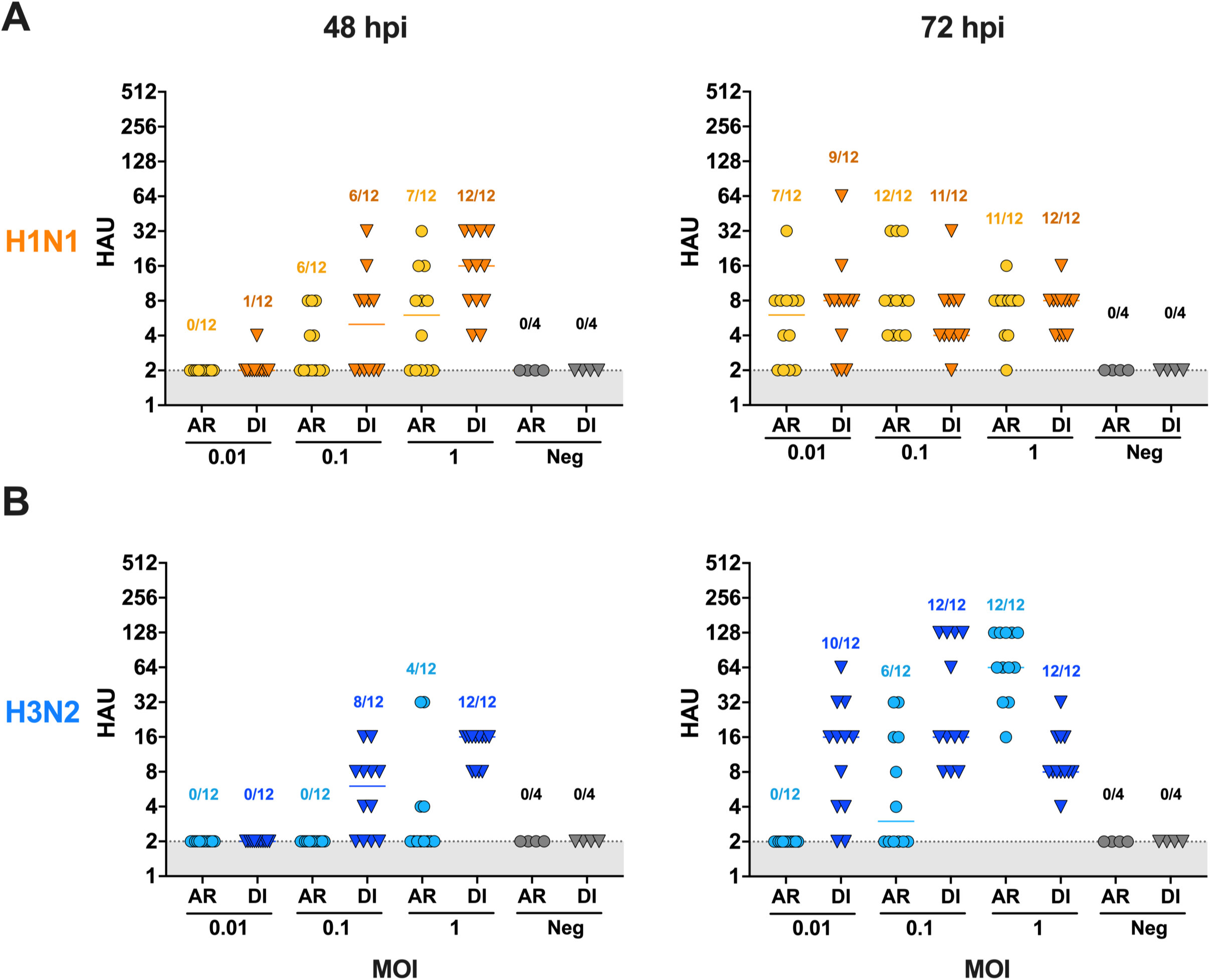
Hemagglutination (HA) assay titration of H1N1 and H3N2 post-aerosol and direct inoculation in differentiated BCi-NS1.1 cells. Differentiated BCi-NS1.1 cells were inoculated via aerosol (AR; circles) or direct (DI; triangles) inoculation of ten-fold dilutions from 1 MOI of **(A)** H1N1 or **(B)** H3N2. Cell culture supernatant was collected at 0-, 12-, 24-, 48-, and 72 hours post-inoculation (hpi). The first three time points (0-, 12-, and 24-hpi) resulted in HAU below the limit of detection and are therefore not shown. Each point represents that HAU from 1 well. Numbers above the points represent the number of wells considered positive/total number of wells. Wells with HAU > 2 were considered positive.

**Figure 7.**
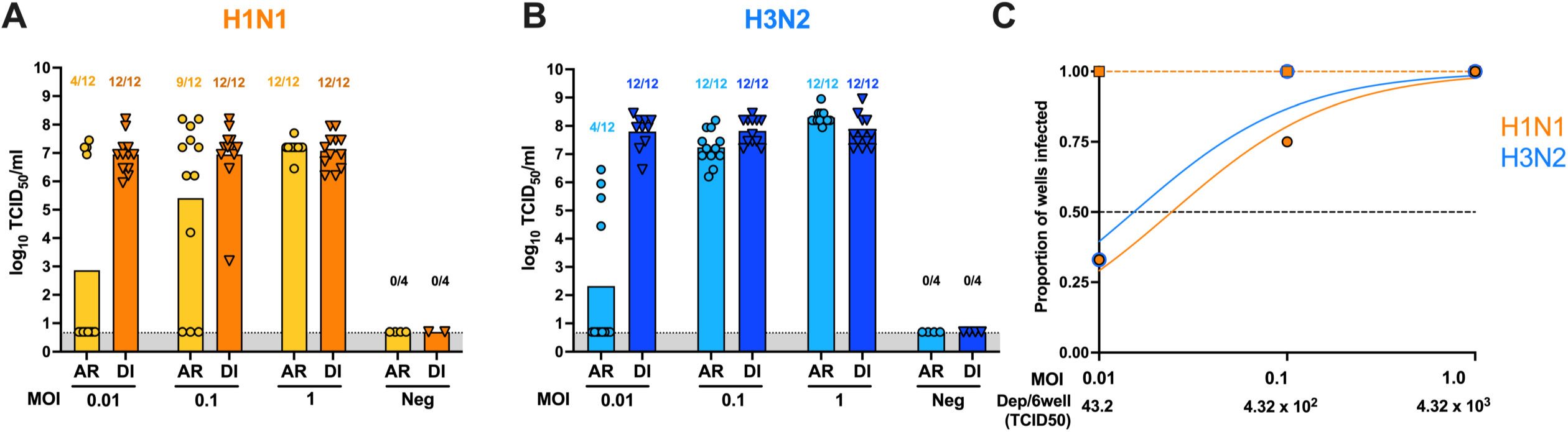
Virus titers of H1N1 and H3N2 post-aerosol and direct inoculation in differentiated BCi-NS1.1 cells. Cultured BCi-NS1.1 cells under air-liquid interface (ALI) conditions for 17 days were exposed to 10-fold dilutions of (**F**) H1N1 or (**G**) H3N2 from 0.01 to 1 MOI via aerosol (AR) or direct inoculation (DI). Cell supernatant was collected at 48 hours post-inoculation (hpi) and infectious virus was quantified and expressed as log10 TCID_50_/mL. Numbers above each bar represent the number of wells considered positive/total number of wells. The proportion of wells with detectable infectious virus observed in **(A)** and **(B)** and their respective dilution factor **(C)** was analyzed in a non-linear regression model to calculate the half maximal effective concentration (EC50). The cumulative deposition of aerosol/6 wells for each inoculum is displayed below. Solid lines with circles indicate aerosol exposure, while dashed lines with squares represent liquid inoculation. Viruses are distinguished by color (H1N1: orange, H3N2: blue)

## DISCUSSION

We generated a calm-aerosol settling chamber system that is accessible and cost-effective to infect a standard cell line (MDCK) and differentiated human airway epithelial cell line (BCi-NS1.1). While previous studies reported exposure systems that aerosolize FLUAV in an *in vitro* and *in vivo* setting (20, 24, 31), these systems generally contain multiple components to control environmental factors and a high cost of implementation and assembly. Therefore, the main objective for designing this system was to encompass a direct layout with minimal components. Prior to aerosol inoculation of different cell types, infectious virus and vRNA collected from two biosamplers were examined to confirm that viable, infectious viral particles were expelled from the nebulizer implemented in this exposure system. Similar quantities of infectious virus from all three viruses were obtained from the SKC BioSampler, suggesting efficient expulsion of infectious aerosols into the aerosol exposure chamber. While this system does not include aerosol measurements and components for environmental manipulation, the system can be utilized for preliminary assessment of aerosol infection efficiency and risk assessment of novel influenza strains.

Using this exposure system, we demonstrate that MDCK cell monolayers can be infected via aerosol by three different subtypes of FLUAV (H1N1, H3N2, and H9N2) using prototypical strains in each case. These virus subtypes were utilized for initial testing because of their relevance in FLUAV transmission, human infection, or potential for zoonosis. Ca04 H1N1 is a prototypical strain isolated from a pediatric patient from the 2009 pandemic, which has been previously demonstrated to transmit via respiratory droplets in ferrets (36, 37). The 2009 pH1N1 lineage can also spillover, reassort, and drift within the swine population, resulting in virus progeny with potential zoonotic transmission (38). The swine-origin Oh/04 H3N2 virus strain used in this study contains the triple-reassortant internal gene (TRIG) cassette, which has been circulating in pigs since 1998 (39, 40). The TRIG cassette includes a human-origin PB1, avian-origin PA, and PB2, and a swine-origin M, NP, and NS gene segments that can support different HA and neuraminidase (NA) combinations of viruses circulating in pigs within North America (39–41). The avian-origin FLUAV strain is an H9N2 subtype, which are zoonotic agents and gene donors to other influenza virus subtypes (H5N1/N6, H7N9, and H10N8/N3) resulting in swine and human infections (42–47). Notably, a previous study showed efficient replication and transmission by direct contact and aerosol of viruses carrying the surface proteins of WF10 but the internal constellation of seasonal H3N2 or pandemic H1N1 viruses in ferrets, suggesting the importance of examining the transmission of H9N2 viruses (48–51). While all three FLUAVs replicated in MDCK cells, the avian-origin H9N2 showed minimal replication in the differentiated BCi-NS1.1 cells via liquid or aerosol inoculation. BCi-NS1.1 cells have been used to examine mechanisms of inflammation (52) and have been reported to be highly similar in morphology, the composition of different cell types, and overall patterns of gene expression compared to primary human airway epithelial cells (53). There are numerous advantages to using the BCi-NS1.1 cells, including extended passage capacity resulting in the opportunity to differentiate larger quantities of cells for extensive experiments, the capacity to utilize engineering techniques for mechanistic studies, and reduced donor-to-donor variation within studies since BCi-NS1.1 cells originate from the same donor (53).While BCi-NS1.1 cells have been exclusively used to examine H1N1 FLUAV infection, specifically the laboratory-adapted A/Puerto Rico/8/1934 (H1N1) and pandemic 2009 H1N1 strains (53, 54), little is known about the presence or distribution of ⍺ 2,3- and ⍺ 2,6-SA. Since WF10 showed minimal replication in the differentiated BCi-NS1.1 cells, screening avian-origin FLUAV in this cell line can be extremely valuable for examining potential zoonosis since H9N2 FLUAVs do not naturally infect humans without further adaptation . Additional studies are needed to confirm susceptibility to H9N2 viruses with these adaptations(55).

Utilizing exposure of MDCK and differentiated BCi-NS1.1 cells to multiple dilutions of each FLUAV, we calculated the AID5_0_ for each virus in each cell type by estimating the cumulative deposition of virus-laden aerosol per 6-well plate exposed. Regarding MDCK exposure, H1N1 (0.021 TCID_50_) had the lowest AID_50_, suggesting that it has a higher efficiency of aerosol infectivity compared to H3N2 (0.13 TCID_50_) and H9N2 (0.035 TCID_50_), especially since all three viruses had similar proportions of wells infected by liquid inoculation. In differentiated BCI-NS1.1 cells, H3N2 (66.2 TCID_50_) had a lower AID_50_ than H1N1 (105.2 TCID_50_), while no detectable HAU was observed after H9N2 exposure. Since numerous studies reported differences in pathogenesis and virus deposition when inoculating animal models via aerosol versus intranasally (17–22), aerosol inoculation of *in vitro* cell culture is valuable in studying molecular markers that could influence the transmission efficiency of FLUAV and virus-cell interactions resembling natural infection in the human respiratory tract.

While the large exposure chamber was initially designed for *in vitro* exposure experiments, several of its components can be adapted for use in an in vivo nose-only aerosol exposure system. Further research is necessary to establish comparability and assess the risk posed by influenza viruses when exposed to HAE cells. The aerosol exposure system presented here successfully infects two cell lines with different subtypes of FLUAV but does not control for various environmental factors that affect aerosol viability and stability, which are crucial for efficient aerosol transmission. Factors like temperature, relative humidity, and UV radiation can significantly impact the survival and transmission of FLUAV aerosols by potentially affecting the stability of proteins, lipids, and vRNA (6). Since the system’s primary objective was to examine the aerosol infectivity of different viruses, direct manipulation of environmental factors was not needed since environmental conditions were relatively similar across experiments. This aerosol exposure system has numerous applications besides assessing aerosol infectivity. The system can also be used to study molecular markers that influence virus-cell interactions resembling natural airborne infection, and aerosol exposure of FLUAV vaccine strains to further improve vaccines with intranasal delivery. In addition to examining FLUAV, this system can be utilized to study other respiratory viruses such as SARS-CoV-2. Using an *in vitro* aerosol exposure system, in conjunction with *in vivo* studies, can aid in expanding the current knowledge on molecular virological markers that influence respiratory virus transmission that can help direct public health preparedness and pandemic risk assessment of circulating FLUAV strains.

## ACKNOWLEDGEMENTS

We thank Lisa Stabler for their assistance in the preparation of HAE cells for histology. We thank the University of Georgia Biomedical Microscopy Core (BMC) for assistance with confocal microscopy.

## AUTHOR CONTRIBUTIONS

DRP developed the original aerosol exposure design. BS and DRP designed the experiments. BS, LCG, FCF, and CJC performed studies and analyzed data. NS and LCM provided guidance and feedback on experimental designs and aerosol calculations. LCM provided edits to the manuscript. MW provided key insights to establish the BCi-NS1-1 cells at the air-liquid interface in the Perez lab and edited the manuscript. AL and DSR consulted and provided feedback on experimental design, data presentation, and edited the manuscript. BS and DRP analyzed the data and wrote and edited the manuscript. All authors read, provided intellectual input and approved the final version of the manuscript.

---

**Supplementary Figure 1. Calculations and exposure chamber parameters to determine the cumulative deposition of virus aerosols per 6 well plate.**

